# Lymphoid origin of intrinsically activated plasmacytoid dendritic cells in mice

**DOI:** 10.1101/310680

**Authors:** Alessandra M. Araujo, Joseph D. Dekker, Kendra Garrison, Zhe Su, Catherine Rhee, Zicheng Hu, Bum-Kyu Lee, Daniel Osorio Hurtado, Jiwon Lee, Vishwanath R. Iyer, Lauren I. R. Ehrlich, George Georgiou, Gregory C. Ippolito, S. Stephen Yi, Haley O. Tucker

**Author notes:** **Corresponding Authors:** Haley O. Tucker, Department of Molecular Biosciences. The University of Texas at Austin, 1 University Station A5000, Austin TX 78712, USA., Joseph D. Dekker, Department of Molecular Biosciences. The University of Texas at Austin, 1 University Station A5000, Austin TX 78712, USA., S. Stephen Yi, Livestrong Cancer Institutes, and Department of Biomedical Engineering, The University of Texas at Austin, Austin TX 78712, USA.

## Abstract

We identified a novel mouse plasmacytoid dendritic cell (pDC) lineage derived from the common lymphoid progenitors (CLPs) that is dependent on expression of *Bcl11a*. These CLP-derived pDCs, which we refer to as “B-pDCs”, have a unique gene expression profile that includes hallmark B cell genes, normally not expressed in conventional pDCs. Despite expressing most classical pDC markers such as SIGLEC-H and PDCA1, B-pDCs lack IFN-α secretion, exhibiting a distinct inflammatory profile. Functionally, B-pDCs induce T cell proliferation more robustly than canonical pDCs following Toll-like receptor 9 (TLR9) engagement. B-pDCs, along with another homogeneous subpopulation of myeloid derived pDCs, display elevated levels of the cell-surface receptor tyrosine kinase AXL, mirroring human AXL^+^ transitional DCs in function and transcriptional profile. Murine B-pDCs therefore represent a phenotypically and functionally distinct CLP-derived DC lineage specialized in T cell activation and previously not described in mice.

## Introduction

Plasmacytoid dendritic cells (pDCs) specialize in the production of type I interferons (IFNs) and promote antiviral immune responses following engagement of pattern recognition receptors. They have been mainly implicated in the pathogenesis of autoimmune diseases that are characterized by a type I IFN signature (notably, IFN-α). Yet, pDCs have also been shown to be able to induce tolerogenic immune responses^5–8^. Due to the clinical significance of pDCs, several studies aimed at mapping pDC lineage derivation have emerged recently. Yet, a complete understanding of pDC development and their existing subsets in mice is still lacking. The transcription factor 4 (TCF4) is required for pDC development and for lineage identity^9–11^. TCF4 is a component of a multiprotein complex that includes both positive and negative regulators^6,12^. One of these components, the transcription factor BCL11a, which is also essential for B cell development^13–15^, induces *Tcf4* transcription and initiates a positive feedback loop with TCF4 to maintain pDC lineage commitment and function^14–16^.

Unlike their conventional dendritic cell (cDC) counterparts, pDCs express transcriptional regulators and markers associated with B-lymphocyte development in addition to BCL11a (e.g. B220, SPIB)^17,18^. These features, along with the established generation of pDCs from myeloid restricted precursors^58,76^, have made it difficult to define pDC lineage affiliation^11,17,19,20^. This has led to the hypothesis that pDC subsets may have distinct origins derived from either the Common Lymphoid Progenitor (CLP) or the Common Myeloid Progenitor (CMP)^18,61^. Beyond the complex nature of their lineage, pDCs with different functional attributes (e.g. variable IFN-α expression levels) or different surface markers (e.g. CD19^+^ pDCs detected in tumor-draining lymph nodes) have also been identified^17,21–23^.

On par with the ever-increasing heterogeneity described within pDC populations, a novel AXL+ dendritic cell (DC) population with many pDC-like properties has recently been discovered in human blood^1,3,4^. While AXL+ DCs express many canonical pDC markers (e.g. CD123, BDCA2/CD303), they also express CD2, the Ig-like lectins SIGLEC1 and SIGLEC6, as well as the activation marker CD81^1^. In a separate study^2^, a similar population of pDCs was shown to express high levels of CD5 and CD81 in mice and humans, two glycoproteins normally associated with the B cell receptor (BCR) signaling complex. The origin of AXL+ transitional DCs is currently unclear and the presence of a homologous population in mice remains muddled. Here, we report the identification of a lymphoid-derived pDC subset (B-pDCs) that shares inflammatory features with myeloid derived *Axl*+ pDC populations in mice. We also demonstrate an in vivo requirement for *Bcl11a* in the transcriptional specification of B-pDCs.

## Results

We previously demonstrated that conditional deletion of *Bcl11a* in the hematopoietic stem cell (HSC) compartment mediated by *Vav-1*-Cre or by inducible *Mx1*-Cre recombinase results in complete abolishment of pDC development^14^. Spurred by previous speculation of pDC origin from the CLP^17,20^, we next selectively deleted floxed (^F^) *Bcl11a* alleles in CD79a^+^ cells ^24,25^, as mediated by *mb1*-Cre *in vivo*. Expression of the *mb1* gene (CD79a) initiates at the LY6D^+^ CLP stage, in B-cell-biased lymphoid progenitors (BLPs)^26^, downstream of LY6D^−^ CLPs (Fig. S1). *Bcl11a^F/F^mb1-Cre* mice (cKO) and littermate controls were examined for pDC frequencies among nucleated cells in the bone marrow (BM). B220^+^ PDCA1^+^ pDCs (all found to be CD11c ^int^) were consistently and significantly reduced by an average of ∼25% (24.8 ± 2.4%) in cKO mice relative to littermates (Fig. 1A-B). Loss of B cells (B220^hi^ PDCA1^-^) served as a gauge of *mb1-Cre* deletion efficiency (Fig. 1A-B). Taken together, these data indicate that a significant proportion of pDCs are derived from BLPs or BLP-derived cells and are *Bcl11a* dependent.

**Figure 1.**
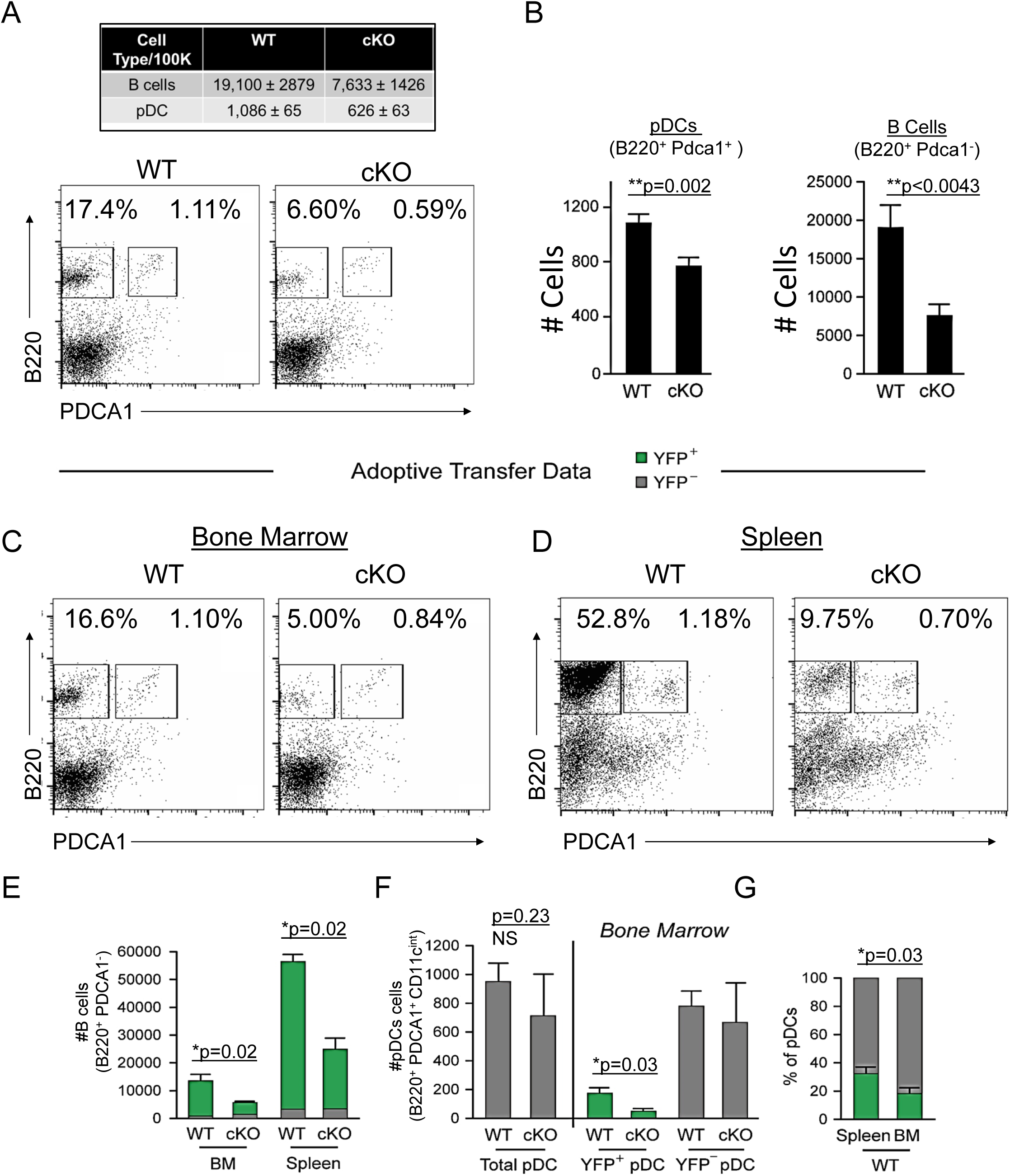
Mb1-Cre deletion of *Bcl11a* identifies a CLP-derived subset of pDCs. (A) Representative FACS plots of pDC (gated as B220^+^ PDCA1^+^) and B cell (gated as B220^+^ PDCA1-) percentages in the bone marrow (BM) of *Bcl11a* ^F/F^ *mb1*-Cre mice (cKO) and littermate controls. (B) Quantification of pDC and B cell populations in the BM of *Bcl11a* ^F/F^ *mb1*-Cre mice (cKO) and littermate controls as cells/100,000 cells. (C-D) Flow cytometric analysis of BM and spleens of recipient mice 8 weeks post BM transplantation. BM was transferred from either BCL11A -sufficient reporter control mice (*mb1*-Cre-YFP) or BCL11A-deficient cKO mice (*Bcl11a* ^F/F^ *mb1*-Cre-YFP) into lethally irradiated C57BL/6J recipients. (E) B cell numbers after BM transplantation in BM and spleens of recipient mice. (F) pDC numbers after BM transplantation in BM and spleens of recipient mice. (G) Comparison of YFP^+^ pDC percentages in the spleen and BM of recipient mice post BM transplantation. Mann-Whitney t-tests were used for all statistical comparisons. Error bars=mean±s.d. The results are representative of two experiments each containing 3-4 mice per group.

To expand these *in vivo* observations, and to confirm that this defect is intrinsic to hematological progenitor cells, we transferred BM from either BCL11A -sufficient reporter control mice (*mb1-Cre-YFP*) or BCL11A-deficient cKO mice (*Bcl11a^F/F^mb1-Cre*-*YFP*) into lethally irradiated wild type C57BL/6J recipients. After 8 weeks, <10% of B cells (B220^+^ PDCA1^-^) in the spleens of *mb1-Cre-YFP* recipients were YFP^−^, confirming elimination of recipient hematopoiesis (Fig. 1D-E). As expected, *Bcl11a^F/F^mb1-Cre*-*YFP* BM resulted in significantly reduced B cell and pDC cellularity compared to *mb1-Cre-YFP* controls, whereas BCL11A-sufficient (YFP^−^) pDCs persisted (Fig. 1C, 1E-F). Approximately 1/3 of pDCs in the spleen of wild-type chimeras were YFP^+^ compared to only 1/5 in BM (Fig. 1G). This increased fraction of YFP^+^ pDCs in the spleen suggests that BLP-derived pDCs preferentially home to that organ. While the residual YFP^+^ pDCs in the BM might be an indication that *Bcl11a* is not shut-off at the most precise developmentally-critical point within the B cell lineage for complete CD79a+ pDC ablation, other hematopoietic lineages were capable of development in normal numbers and contained a paucity of YFP^+^ cells (Fig. S2). Of note, BCL11A-deficient progenitors yielded higher splenic T cell chimerism at the expense of B cells and pDCs, but less than 2% of T cells were YFP^+^ (S2, and not shown). This indicated that *mb1-cre* expression occurs subsequent to T-B lineage divergence, consonant with previous observations^26,68,69^ and our *mb1-cre* progenitor analysis in which YFP^+^ cells are confined to the BLP derived compartment (Fig. S1). Because of their exclusive lymphoid derivation post T-B bifurcation, we were prompted hereafter to refer to this pDC lineage as “B-pDC”.

Relative to CD79a^-^ pDCs, resting B-pDCs express higher levels of MHC Class II (Fig 2A-B) suggesting that they may be primed for immediate response to pro-inflammatory signals. To test this hypothesis, we delivered TLR9 ligand (CpG:ODN) into *mb1-Cre*-*YFP* mice via tail vein injection, and splenic pDCs were phenotyped via flow cytometry 24 hours later. While both pDC and B-pDC compartments expanded relative to controls (Fig. 2C-D), the YFP^+^ B-pDC fraction increased almost 2-fold above that of the YFP^−^ pDC fraction in numbers (Fig. 2E). Additionally, fold increase of B-pDCs coexpressing CD83 and CD86 upon activation was also significantly higher than that of YFP^-^ pDCs (Fig. 2F-G). These results suggested that relative to YFP^-^ pDCs, B-pDCs are intrinsically activated and primed for rapid expansion upon TLR9 engagement. To confirm the functional phenotype of B-pDCs, we tested their ability to secrete cytokines known to be elicited by pDCs after Toll-like receptor (TLR) engagement. Specifically, we tested each pDC lineage for the production of IFN-α or IL-12p40 when activated by TLR9-bound CpG oligonucleotides. We sorted B-pDCs and pDCs, engaged TLR9 with CpG:ODNs for 24 hours and collected supernatant for cytokine specific ELISAs. IFN-α production was almost negligible in B-pDC (Fig. 2H), yet IL-12p40 production was significantly augmented over pDC (Fig. 2H). To test their ability to expand T lymphocytes in culture, sorted B-pDCs and pDCs were incubated with CpG:ODNs and co-cultured with freshly isolated CFSE-labeled lymphocytes. After 6 days, co-cultures were stained for CD3 and CFSE-negative cell percentages were recorded (Fig. 2I-J). Altogether, our results showed B-pDCs were significantly better at expanding T cells in co-cultures than conventional pDCs (Fig. 2I-J) upon TLR9 activation.

**Figure 2.**
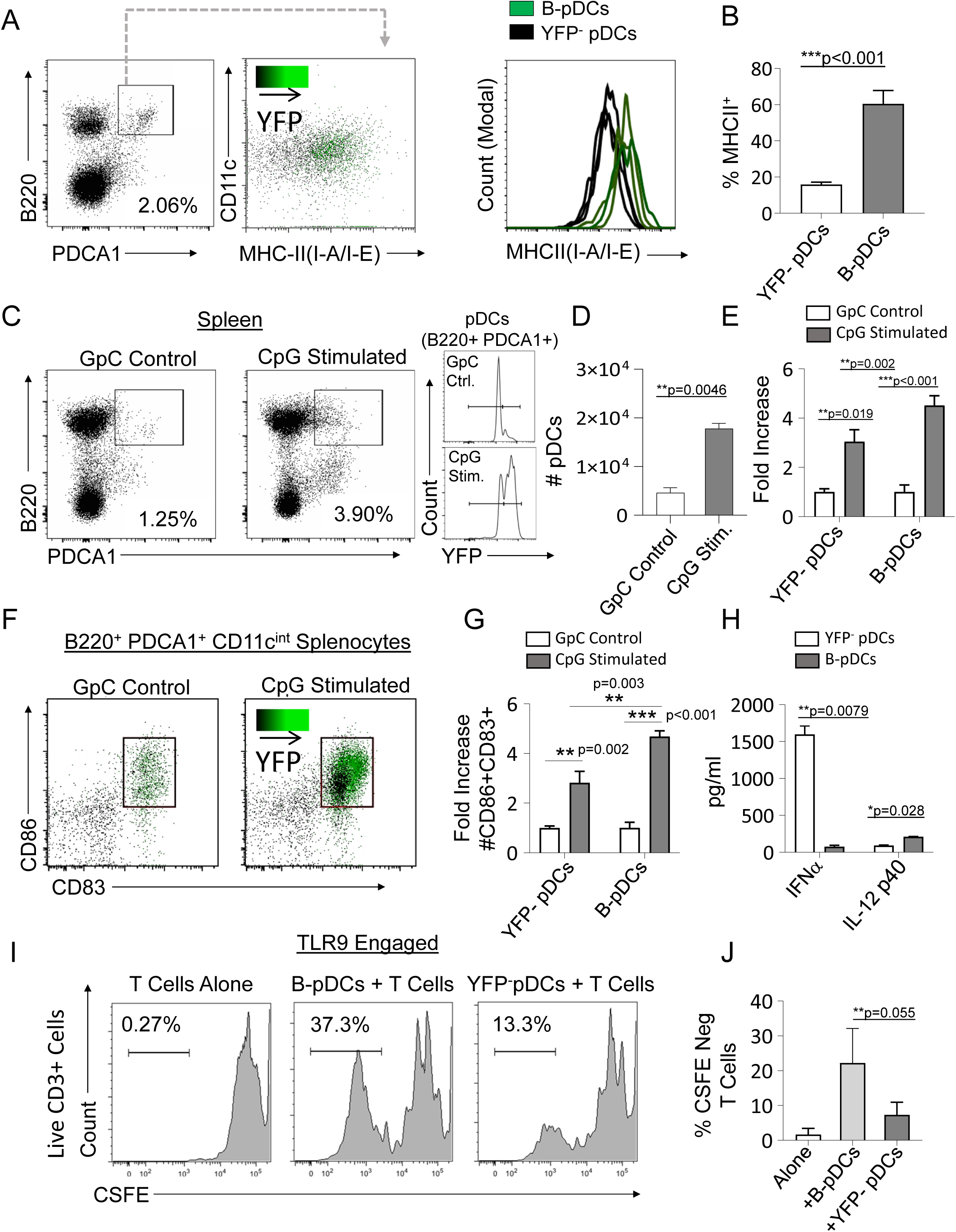
CLP-derived B-pDC are licensed for T cell activation. (A-B) Comparison of MHCII MFI in YF^+^Pand YFP^-^ pDCs from *Mb1*-Cre-YFP reporter mice via flow cytometric analysis. (C-D) *Mb1*-Cre-YFP mice were injected with 50 ug/ml (100 ul) of CpG:ODN or control GpC:ODN and analyzed via flow cytometry for splenic pDC numbers/100,000 cells. (E) Flow cytometry quantification of YFP^-^ pDC and B- pDCs (YFP^+^) fold change in cell numbers upon CpG:ODN in vivo challenge. (F-G) Fold change in CD86^+^CD89^+^ cell numbers for each pDC population in mice restimulated with CpG:ODN. Difference in CD86^+^CD89^+^ cell numbers between stimulated B-pDCs and YFP ^-^ pDCs was also significant. (H) In vitro TLR9 engagement of B-pDC or pDC for ELISA against IFN-α orIL-12p40. (I) T cells were magnetically isolated from wildtype C57BL/6J mouse splenocytes using MACs columns, labelled with CSFE, and then cultured (2.5x10^4^/well) alone or with CpG:ODN activated B- pDCs or pDCs (5x10^3^/well) for 6 d. (J) The percentage of CFSE-negative CD3^+^ T cells in co-cultures were significantly higher in B-pDC compared to pDCs. Mann-Whitney t-tests were used for all statistical comparisons. Error bars=mean ±s.d. The results are representative of three independent experiments each containing at least four mice per group.

To further elucidate their phenotype at the genetic level, we performed RNA-seq analyses of purified B-pDCs and compared them to classical, myeloid-derived bulk pDCs. Expression of hallmark pDC genes was confirmed by RT-PCR of shared pDC markers (Fig. S3A). While the overall gene expression patterns were highly similar across the two subsets, ∼1% of transcripts (∼220 genes) differed significantly (*q* value< 0.05) (Fig. S3B). Among the top overexpressed genes in B-pDCs were *Lyz1, Ccr3, Cd86, Id2, Axl, Siglec1, and Cd81* (Fig. S3C). Differentially expressed transcripts generated Gene Ontology (GO)^27^ or Panther^28^ terms including “immune response”, “inflammatory response”, “cell activation”, and “regulation of immune response” (p=1.36x10^-17^, 2.15x10^-12^, 3.67x10^-11^, and 1.45x10^-10^, respectively (Fig.S3C). Gene set enrichment analysis (GSEA) revealed elevation of each of these same GO/Panther terms within the B-pDC subset as compared to a pDC-related GSEA control dataset that showed no enrichment. Next, we compared all genes expressed by both pDC populations to one another and to published^29^ RNA-seq of BM-derived mouse pre-B cells (B220^+^ IgM^-^ Kit^-^ CD25^+^)—a post-CLP B cell progenitor (pre-B) population. As shown in Figure S3D, pDCs and B-pDCs expression levels were strongly correlated with one another relative to early B cells (R^2^ values = 0.8959, 0.4145 and 0.404, respectively) (Fig.S3D). Collectively, these data support our contention that B-pDCs are functionally distinct from classical pDCs and specialize in inflammatory responses, antigen presentation, and T cell activation.

AXL^+^ transitional DCs (tDCs or AS-DCs)^1,57^ have been identified in humans by the high expression levels of the genes *AXL*, *CD81*, *CD86*, *LYZ2*, *C1QA*, *CD2*, and *C1QB* relative to classical pDCs ^1–4^. They also exhibit many classical pDC features, having previously been described phenotypically as a “continuum” between pDC and cDC2 populations ^2^. In order to confirm AXL^+^ pDCs are also present in mice and corroborate our RNA-seq findings, we phenotyped B-pDCs through imaging flow cytometry. First, we again verified that our primary gating scheme effectively encompassed bona-fide pDCs using imaging flow cytometry. As expected, B220^+^ PDCA1^-^ cells were >95% CD19^+^ SIGLEC-H^-^ (B cells). Most importantly,>95% of B220^+^ PDCA1 ^hi^ cells were SIGLEC-H ^+^ and CD19^-^, ruling out any pDC population contamination (Fig. S.4A-B). At the protein level, we were able to stain mouse BM cells with anti-AXL antibodies without issues and confirmed that while ∼60% of B-pDCs were positive for AXL, only ∼10% of YFP negative pDCs expressed this protein (Fig. S4A, C-E). Finally, imaging of BM cells showed that B-pDCs resembled classical secretory pDCs morphologically, presenting a large nucleus and defined plasma-like features, in contrast to B cells (Fig. S4A). This preliminary phenotypic analysis led us to speculate that similarly to human AXL^+^ transitional DCs, B-pDC might also prioritize their secretory capacity for copious transcription and secretion of alternative immune system modulators. Thus, we performed high resolution 10x single cell RNA-seq analyses of magnetically sorted PDCA1^+^ bone marrow cells (96% purity as determined by flow cytometry) and compared the B-pDC transcriptional profile to that of B cells and other pDCs. Principal component-based clustering and UMAP visualization of sorted PDCA^+^ cells yielded 22 discrete clusters. To identify cell populations in an unbiased manner, we uploaded the top 1000 differentially expressed genes (DEGs) from each cluster into CIPR (Cluster Identity Predictor)^66^, which uses the reference database ImmGen (mouse) to calculate cell identity scores (S. Table 1). Seurat clusters were named according to the consensus of the top 5 cell ID hits generated by CIPR (Fig. 3A). We confirmed CIPR assignments by evaluating cluster DEGs and expression of markers associated with specific myeloid cell populations (Fig. S5). PDCA1^+^ cells comprised 9 distinct pDC populations (including B-pDCs, which were unanimously classified as pDCs by CIPR identity scores despite its unique *Cd79a* expression). Of note, pDC8 had one of the lowest ID scores among consensus pDCs (S. Table 1) and yielded CIPR identity hits for both pDCs and common myeloid progenitors “CMPs” within its top 5 ID hits (not shown). This suggested that this single cluster (1.7% of total cells) represented a mixed population of pDCs and differentiating CMPs expressing high levels of *Hbb* family genes (hemoglobin associated genes) under the chosen default Seurat clustering resolution settings. Because of pDC8 mixed identity, we excluded this cluster from our further single cell transcriptional analysis. Besides pDCs, a small B cell cluster (2.75% of all cells analyzed) expressing low levels of *Bst2* (PDCA1 gene), as well as distinct subpopulations of macrophages (Mac1, 2, and 3), monocytes (Mono1, 2 and 3), and granulocytes (Granul1, 2, and 3) were also identified. Two stem-progenitor populations (Prog1 and 2) were present in our cell pool (1.12% and 2.86% of cells analyzed, respectively) (Fig. 3A-B). The Prog2 population expressed high levels of genes associated with pDCs, including *Siglech*, *Bst2*, *Flt3, TCF4,* and *Irf8* and had a “CDC” (common dendritic cell progenitor) signature according to its CIPR reference ID (Fig. S5, S. Table 2). In addition, the majority of cells in the Prog2 cluster were *Csf1r*^-^, *Il7r*^+^, *Siglech*^+^, *Ly6d*^+^ (Fig. S5, S. Table 2), resembling the pro-pDC myeloid precursor population described by Rodriguez *et al*, as well as Dress and colleagues^67,70^. In contrast, Prog1, which clustered near B cells and was identified as “MLP” (multi-lymphoid progenitors entering the CLP stage), expressed several pre/pro B cell genes, including *Jchain, Iglc1, Iglc2, Igha, Igkc* (S. Table 2). Direct DEG comparison between Prog1 and Prog2 clusters confirmed Prog1 cells exhibit a more “Pre/Pro B cell-like” developmental profile and low expression of markers that are “pDC biased” relative to Prog2 (S. Table 2). After clustering and profiling PDCA1*^+^* cells, we subsetted the data in order to compare B-pDCs gene expression to only B cell and pDC clusters using Wilcoxon rank sum tests. We plotted several genes commonly associated with B cells, and pDCs. Except for *Cd79a (Mb1)* and *Cd79b*, B-pDCs showed negligible expression of mature B cell associated genes such as *Pax5*, *Cd19* and *Cd22* (Fig. 3C). Expression of canonical pDC genes was present but reduced in B-pDCs relative to classical pDC subsets (Fig. 3D). Our analysis highlights the B-pDC cluster uniqueness among pDC subsets and further supported its lineage divergence from mature B cells.

**Figure 3.**
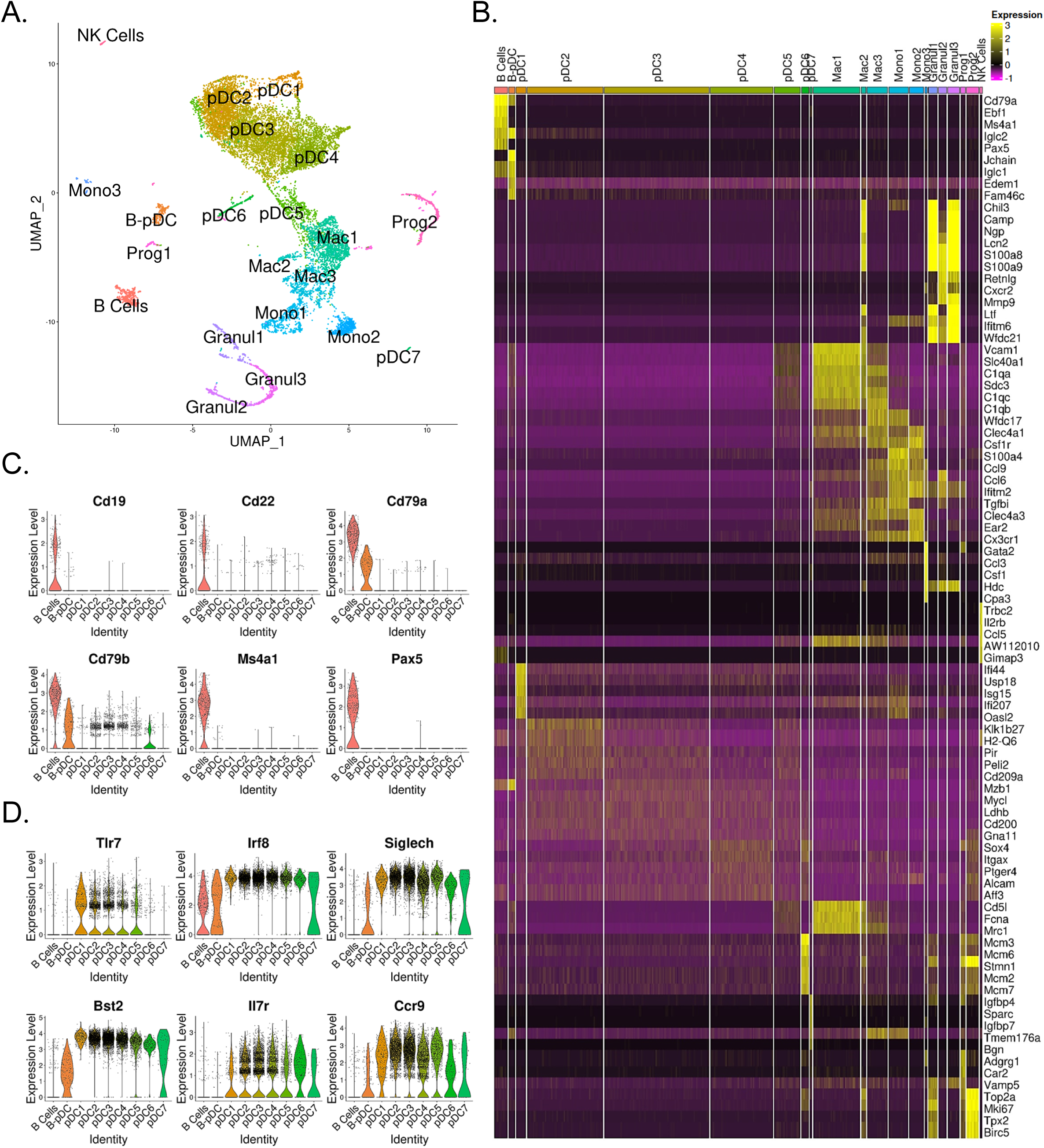
Single cell RNA-seq analysis of mouse PDCA1^+^ bone marrow cells. (A) UMAP generated by Seurat clustering analysis. Cluster identities were assigned using the top 5 consensus CIPR identity scores. (B) Heatmap of the top 5 DEGs from each cluster. After Seurat clustering, cell reads were subsetted to include only cells classified as B cells, pDCs, and B-pDCs. DEGS were generated for the new data subset and markers associated with (C) mature B cells (D) classical pDCs were plotted as violin plots.

In order to temporally examine lymphoid transcriptional commitment to B cell or B-pDC lineages, we inferred developmental trajectories for B and B-pDC single cells. Pseudotime is a measure of how much progress an individual cell has made through a process such as differentiation^59,60^. In other words, cells that belong to a single cluster can be segregated across various transcriptional “states” based on number of reads per cell of a particular set of developmental genes. Although we cannot say the pre/pro B cell-like Prog1 cluster represents an immediately adjacent common progenitor for B cells and B-pDCs, we confirmed B-pDCs are not a direct precursor for B cells based on their single cell developmental trajectory. Cells in the Prog1 cluster were chosen as state “0” based on their abundant expression of lymphoid progenitor associated genes (Fig. 4A). Analysis of uniquely expressed genes showed Prog1 downregulated several multipotency markers such as *Stmn1*, *Gata2*, and *Ctla2a*, while B cells increased expression of transcription factors needed for B cell commitment (e.g., *Pax5*, *Ebf1*, and *Ms4a1*) (Fig. 4B and 4C). Although B-pDCs did not express detectable levels of most B cell commitment markers evaluated, they did retain expression of early B cell receptor associated genes, including *Ly6d*, *Ighm*, *Igkc*, *Iglc2*, and *J Chain* (Fig. S6A). Because these early B cell related markers are also expressed in plasma cells, we performed flow cytometric analysis of B220^+^ PDCA1^+^: CD79a^+^ cells. We discovered that B-pDCs express negligible levels of TACI (a transcription factor that silences the B-cell developmental program ^62^), and that expression of CD138 and IgG1 was also negligible (Fig. S6B, and not shown), thus ruling out any major plasma cell overlap at the protein level. Lineage analysis also revealed that B-pDCs enhanced expression of the pDC associated markers *Siglech*, *Tcf4*, *Bst2*, and also of *Xbp1* (a marker expressed at high levels in secretory cells). Committed B cells in turn downregulated these markers as they progressed through pseudotime (Fig. 4D). In summary, lineage analyses revealed that as B-pDCs develop, they retain expression of both early B cell receptor genes and pDC associated genes, diverging from B cells through continued suppression of transcription factors required for terminal B cell differentiation.

**Figure 4.**
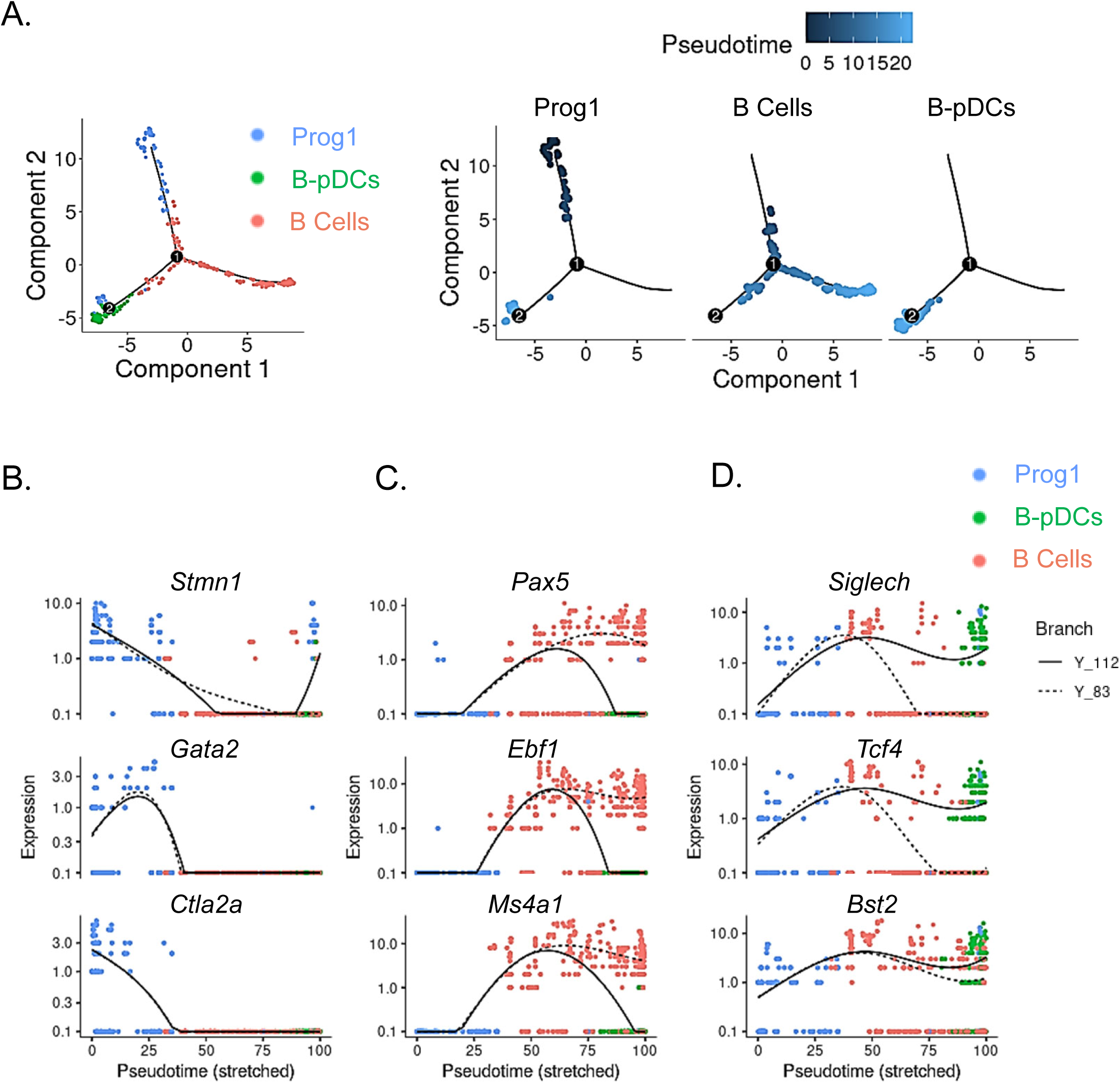
Single cell trajectory analysis of Prog1, B cells, and B-pDC clusters. (A) Single cells belonging to clusters Prog1, B cells, and B-pDCs were ordered and plotted as a function of pseudotime based on uniquely expressed markers using the unsurpevised Monocle dpFeature. Cluster Prog1 was classified as the root of the trajectory given its high expression of pre-pro B cell and stem cell associated markers. DEGs of interest were plotted as a function of pseudotime in Prog1 (B), B cells (C), or pDCs (D) using Monocle. Branch Y_112 represents the expression kinetic trend of the B-pDC cluster (green), while branch Y_83 represents the expression kinetic trend in expression of B cells according to branched expression analysis modeling, or BEAM.

In order to determine if murine B-pDCs are homologous with AXL+ human DCs, we subsetted pDCs clusters and plotted their relative expression of AXL+ DC associated genes. We identified B-pDCs and pDC5 as “noncanonical pDCs” due to their distinguishing high expression of *Axl* (Fig. 5A). Accordingly, *Axl* negative pDCs are referred hereafter as “classical pDCs”. We then analyzed the top DEGs shared between B-pDCs and pDC5s using the function “FindConservedMarkers”. *Lyz2*, the complement genes *C1qa, C1qb, C1qc*, and *Cd5l* were the top conserved markers for these two pDC subsets (Fig. 5A). Of note, *Cd5l*, a macrophage associated secreted protein, has been shown to play a major role in initiating inflammation and maintaining humoral responses. ^71–73^ Unexpectedly, the number of reads for myeloid markers previously used to define human AXL+ DCs via CyTOF or RNA-seq^1,57,67,69,70^ was negligible in murine pDCs and weakly correlated with expression of *Axl* itself in mice (Fig. S7). *Cd81* expression, however, was highly expressed in B-pDCs (Fig.S7A). Of note, *Cd81* and *C1qb* have been described as hallmark genes of a discrete mouse pDC population that does not produce type I IFN and yet mediates important immune functions previously attributed to all pDCs^2,37^. Altogether, our analysis revealed that constitutive expression of *Axl* and innate activation markers are present in more than one transcriptionally distinct murine pDC profile, stressing the functional heterogeneity and plasticity of murine pDCs.

**Figure 5.**
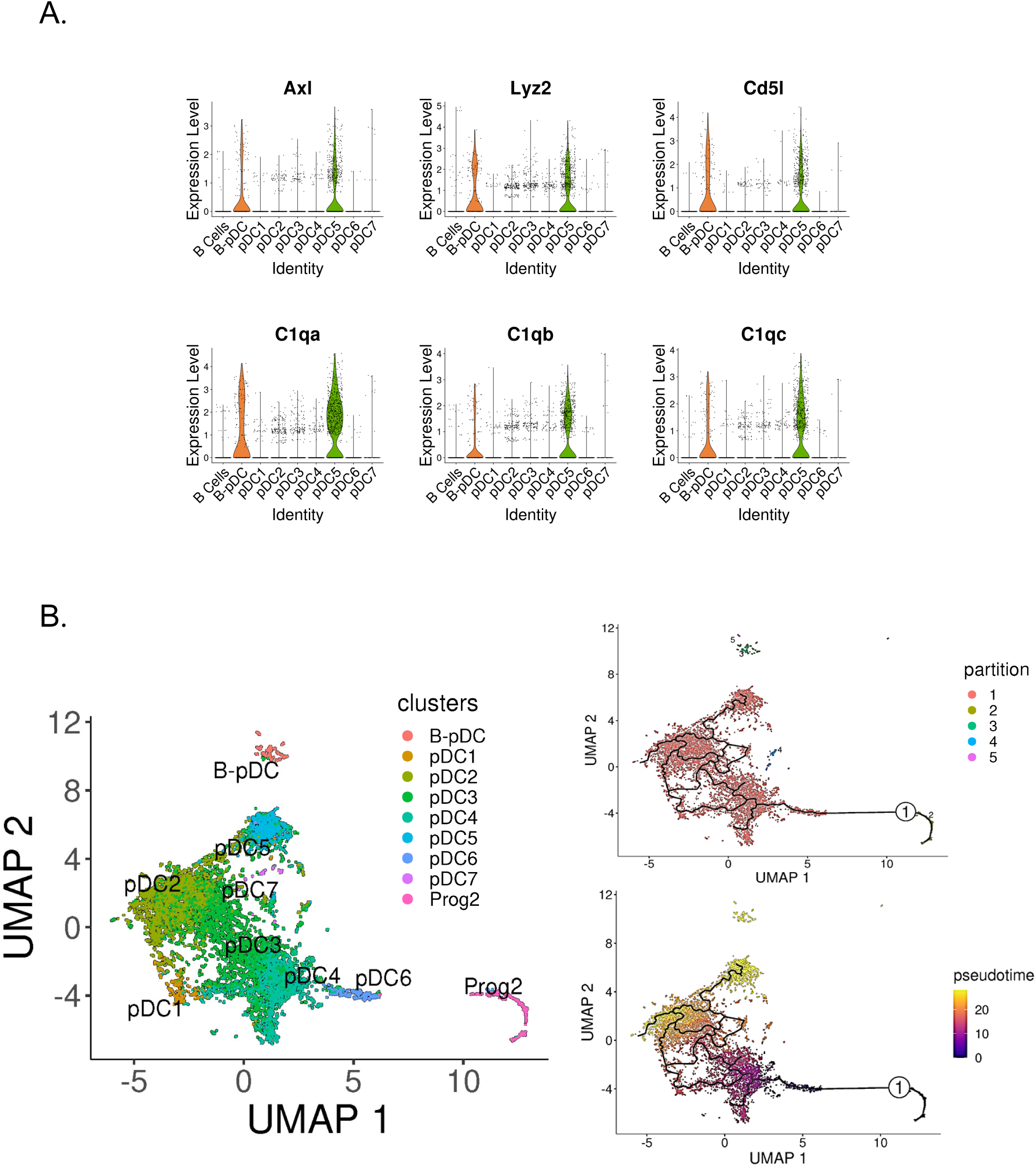
Identification of Axl^+^ non-canonical pDCs in mice. (A) Violin plots depicting relative expression of conserved “non canonical *AXL*^+^ DCs” associated markers in mouse pDC clusters. (B) BM scRNA-seq data was subsetted to include only pDCs and Prog2 subsets. We used Monocle3 for partition-based learning of cell developmental trajectories. Louvain partitions were generated from single cell reads and their developmental trajectory was inferred using SimplePPT (top right quadrant). Pseudotime across clusters is shown in the (lower bottom quadrant).

Noncanonical AXL+ DCs in humans have been shown to be derived from a myeloid pro-pDC progenitor^67,69,70^. In mice, Feng and colleagues used clonal lineage to show that pDCs and cDCs originate from a common Flt3-dependent pathway of differentiation originating from transient CX3CR1^+^ progenitors. Since the authors state that their finding does “not completely rule out the common origin of pDCs and B cells, which may bifurcate earlier in hematopoiesis”, we set out to corroborate the hematopoietic origins of pDC5 and B-pDCs. We first calculated pDC5’s “community connectedness” ^74^ to other pDC clusters and plotted their developmental trajectories originating from the pro-pDC progenitor cluster Prog2 using Monocle3^75^. Partitioning Prog2 and pDC single cells into Louvain communities yielded 5 supergroups, with the largest group encompassing most pDC clusters, including pDC5. The other groups consisted of non pDCs (Prog2 cells), B-pDCs, and a few single cells (>20 cells) genotypically distinct from their original pDC clusters (Fig. 5B). Ultimately, this statistical grouping confirmed that B-pDCs were not developmentally linked to “classical” AXL negative pDCs or *Axl*+ pDC5 cells. Accordingly, B-pDCs did not integrate into the predicted lineage trajectory originating from Prog2, whereas the *Axl+* pDC5 population did (Fig. 5B). Both B-pDCs and pDC5s exhibited a similar late developmental pseudotime, while clusters pDC6 and pDC4 were the first to develop from Prog2. Collectively, our data establish that murine non canonical pDCs (AXL+) consist of at least two major pDC subpopulations, which develop from lymphoid and myeloid progenitors, respectively.

## Discussion

Dendritic cell–subset biology, development, and the ensuing nomenclature have long been unclear and everchanging. Here, we provide definitive evidence in support of the long-suspected “lymphoid past” of pDCs by establishing their ability to arise *in vivo* from CLP derived progenitors with B/pDC bipotential lineage capacity^17–19,43,44^. Our data shows that the murine pDC compartment is bipartite, being comprised of B-pDCs—diverted from the CLP post T-B bifurcation—as well as myeloid-derived classical pDCs. Unlike most myeloid derived pDCs, B-pDCs constitutively express high levels of multiple innate activation markers, including *C1qa, C1qb, Lyz2, and Cd81.* Functionally, they expand more readily after TLR9 engagement than classical pDCs (either through increased proliferation or differentiation of other cell types) and excel at activating T cells in culture. While further functional definition awaits discovery, our work provides a framework for the identification and segregation of the B-pDC lineage (comprising almost 1/5 of the total pDC compartment) from other myeloid-derived pDC subpopulations. Primarily, our observations support the hypothesis that DC functionality derives primarily from ontogeny rather than from tissue environment ^50^, exemplified by evolution of a specialized pDC lineage from a lymphoid progenitor. In the case of the B-pDCs, our data suggest such a cell may deviate from B cell commitment after *Ly6d* expression, thought to be the earliest marker of B cell specification^26^.

Most noticeably, our newly described B-pDC population expresses high levels of the tyrosine kinase AXL and specializes in T cell activation through displaying high levels of MHC II and costimulatory markers. Of note, ablation of AXL in mice has previously been shown to increase expression of type I IFN while impairing IL-1β production and T cell activation during viral infections. ^63,64^ Our related findings that AXL+ B-pDCs excel at T cell priming but exhibit reduced IFNα expression might suggest these DCs play an important role advancing T cell expansion, while curbing innate immune responses that can result in autoimmune damage. Of note, whether induction of IFN-I production i*n vivo* could also affect CLP and increase the amount of YFP+ lymphoid progenitors and thus B-pDC output is unclear. Further research is required to answer this question.

In addition to describing a novel B cell-like pDC population, we profiled murine can develop in parallel from either myeloid or lymphoid progenitors, with lymphoid derived B-pDCs retaining expression of additional early B cell genes that are not expressed in myeloid derived pDCs. Intriguingly, ChIP-seq analysis done by our lab of BCL11A target binding in multiple human cell lines suggests that an evolutionarily conserved transcriptional hierarchy might distinguish AXL^+^ pDCs and B cells in humans as well (unpublished data). The demarcation of lymphoid derived B-pDCs as one of the AXL^+^ pDC populations found in mice may help clarify the perceived plasticity of the pDC compartment in normal and disease contexts^51^, as well as provide a new cell for targeted study within the context of autoimmune disease, cancer, and infection models.

## Supplemental Figure Legends

**Supplemental Figure 1.**
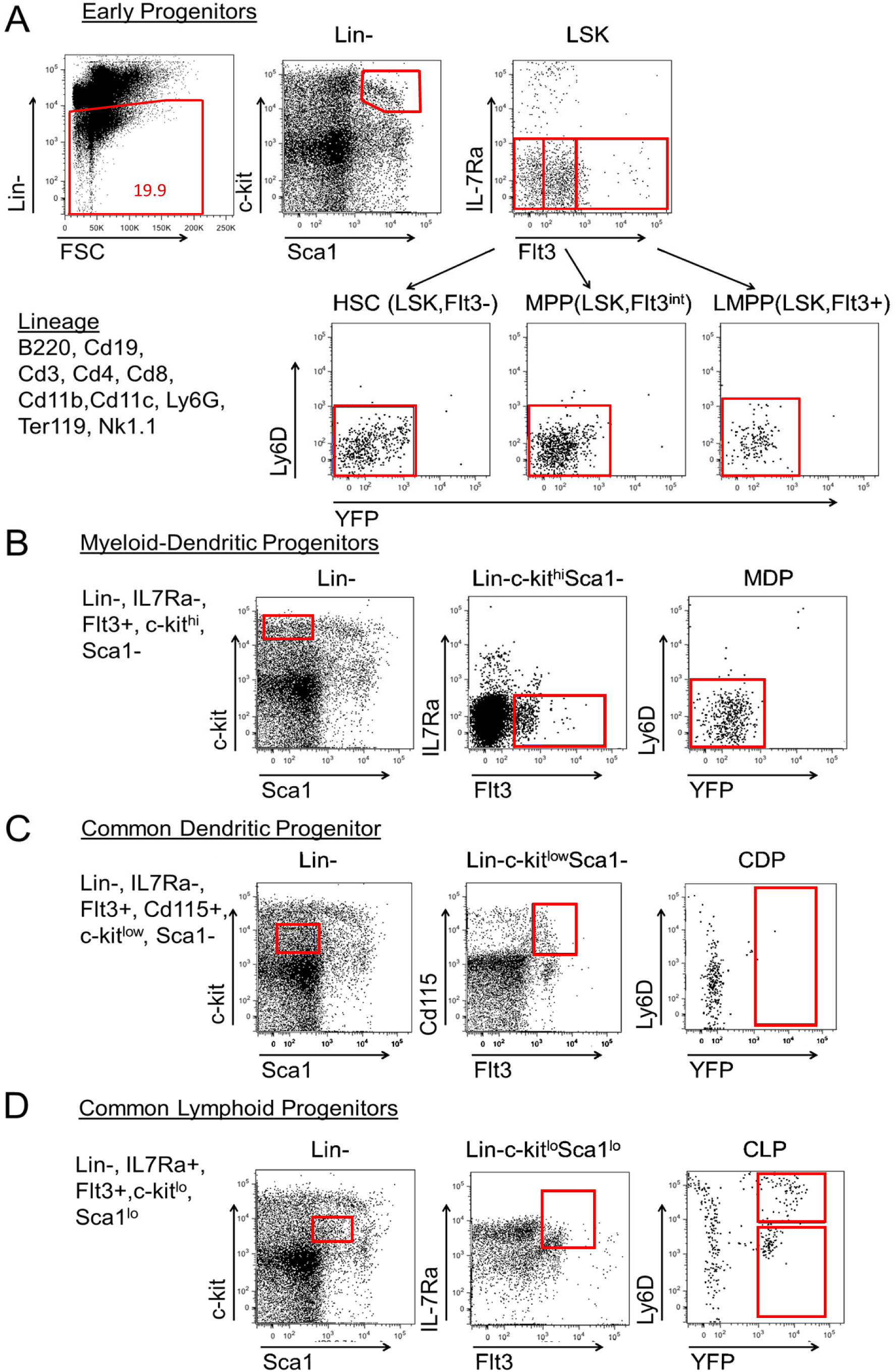
Progenitor population analysis for mb1-cre driven YFP expression. We analyzed YFP expression in BM progenitors to determine when and where mb1-Cre is active (representative plots shown). (A) Lineage negative cells were analyzed to examine LSK Hematopoietic progenitor (Lin^-^, Sca-1^+^, c-Kit^+^), MPP (LSK, Flt3^int^), and LMPP (LSK, Flt3^hi^), populations. Virtually all cells were YFP^-^. (B) Myeloid Dendritic Progenitors (MDP; Lin^-^, IL7Rα^-^, Flt3^+^, c-kit^hi^, Sca1^-^) and (C) Common Dendritic Cell Progenitors (CDP; Lin-, IL7Ra-, Flt3^+^, c-kitlo, Sca1-, CD115^+^) were also negative for YFP expression, while (D) Common Lymphoid Progenitors (CLP; Lin^-^, IL7Ra^+^, Flt3^+^, c-kit^lo^, Sca1^lo^) contained YFP^+^ cells. The results are representative of three independent experiments each containing at least three mice per group.

**Supplemental Figure 2.**
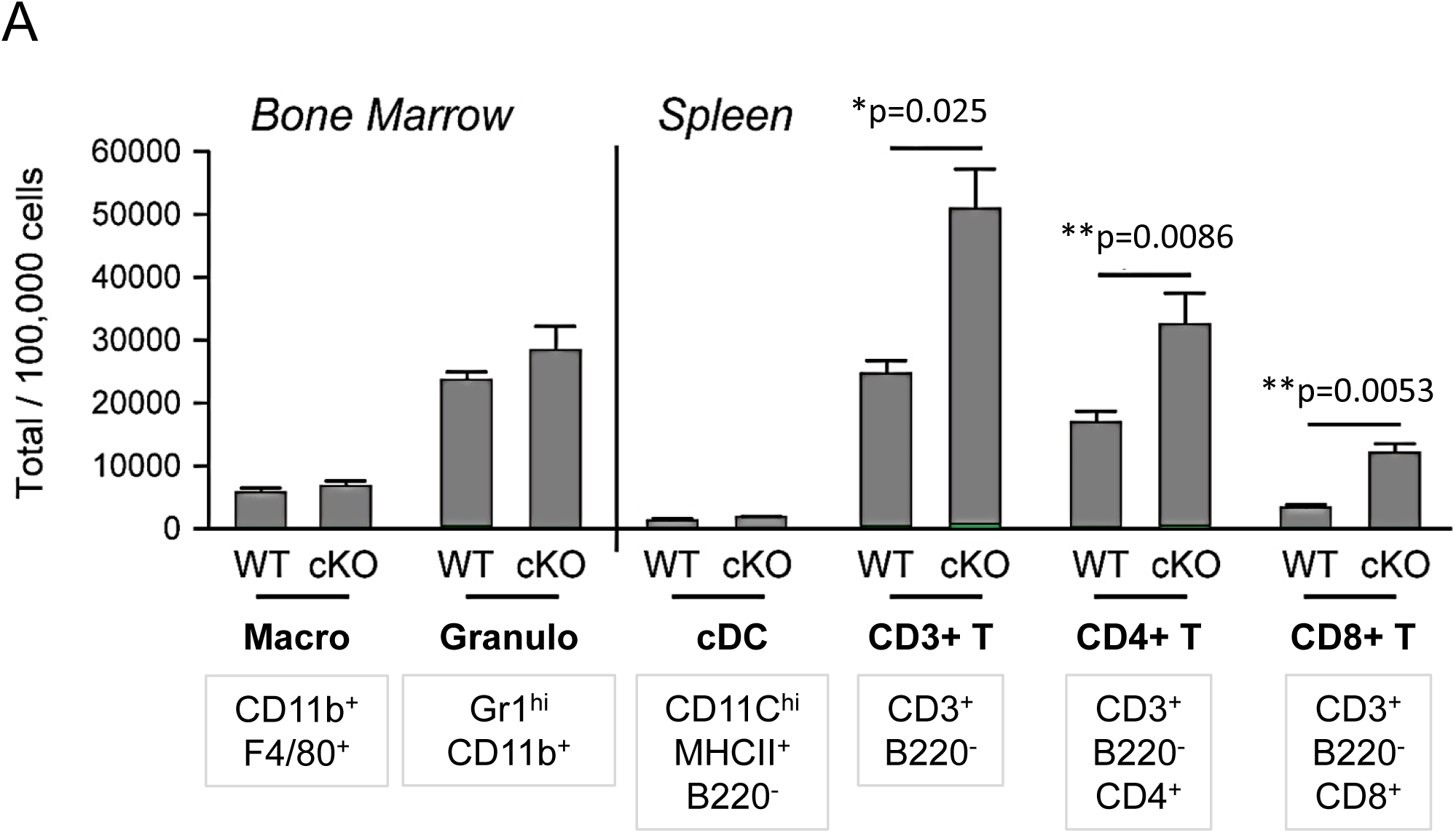
Cell distributions in the spleen of adoptively transferred recipient mice. Both *Bcl11a* ^F/F^ *mb1*-Cre^+^ and *Mb1*-Cre-YFP adoptively transferred recipient mice reconstituted other splenic cell types in normal numbers, including BM macrophages (CD11b^+^ F4/80^+^), granulocytes (Gr-1^+^ CD11b^+^) and splenic cDCs (CD11c^+^ CD11b^+^ B220^-^). Total T cells (CD3^+^ B220^-^) as well as CD4^+^ and CD8^+^ subsets were significantly increased in number in proportion to B/pDC cell loss. Mann-Whitney t-tests were used for all statistical comparisons. The results are representative of two independent experiments each containing 3-4 mice per experimental group. Error bars=mean±s.d

**Supplemental Figure 3.**
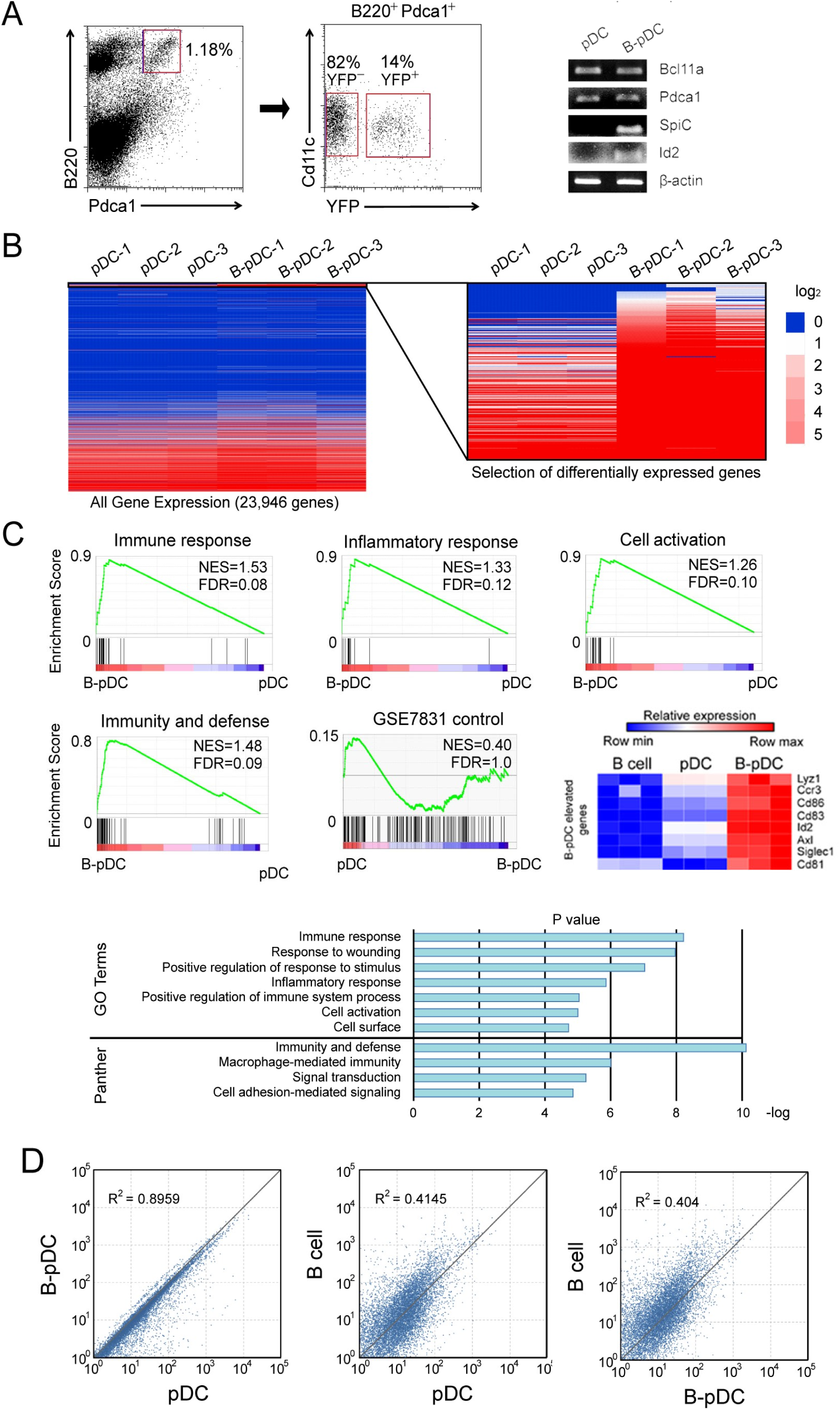
Transcriptional analysis identifies two populations of pDC in mice: myeloid-derived classical pDC and CLP-derived B-pDC. (A) Bone marrow pDC (B220^+^ PDCA1^+^ CD11c^int^ CD11b^-^) were sorted based on expression of YFP. Four mice were pooled for each isolated RNA sample, for a total of three pDC and three B-pDC groups from 12 mice. RT-PCR was used to confirm that *Bcl11a, Bst2, SpiC, and Id2* expression match RNA-seq trends between pDC and B-pDC (B) RNA-seq was performed for gene expression analysis of pDC vs B-pDC and 220/23,946 genes (∼1%, left) were significantly differentially expressed (q value < 0.05, right heatmap, Log2 expression difference displayed). (C) Gene Set Enrichment Analysis (GSEA). Normalized enrichment score (NES) and false discovery rate q-values(FDR);FDR≤.25isconsideredsignificant^56^. GO Term or Panther-derived pathways identified by DAVID analysis of ∼220 differentially expressed genes between the pDC and B-pDC subsets^28,29^. Among the top B-pDC DEGs were *Lyz1, Ccr3, Cd86, Cd83, Id2, Axl, Siglec1, and Cd81.* (D) Scatter plot comparisons of all genes with Reads Per Kilobase of transcript, per Million mapped reads (RPKM) >1. Correlations of pDC vs B-pDC (R2 value = 0.8959), B cell vs. pDC R2 value = 0.4145) and B cell vs. B-pDC (R2 value = 0.404) are indicated.

**Supplemental Figure 4.**
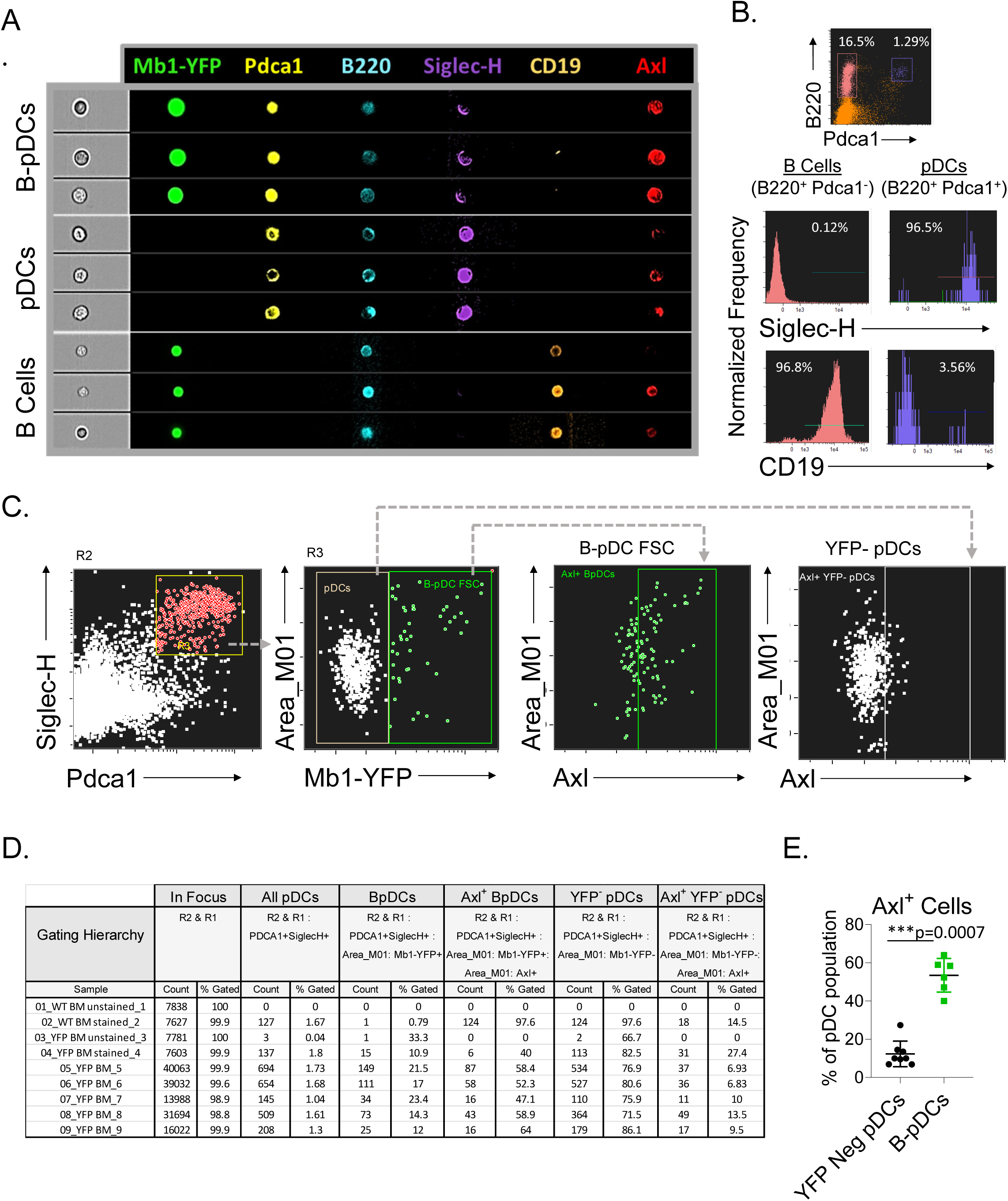
Imaging flow cytometry as a tool to quantify AXL expression in pDC populations. (A-B) Representative Imaging flow cytometry of manuscript gating scheme used to define B cells and pDCs and histograms for SIGLEC-H, AXL and CD19 in B cells and pDCs (C) Representative Imaging flow cytometry plots of AXL^+^ pDCs (here alternatively gated as SIGLEC-H^+^ PDCA1^+^) in *Mb1*-Cre-YFP reporter mice bone marrow cells (n=5). (D) Complete statistics table generated with IDEAS for Image Stream analysis including non-stained wildtype controls. (E) Comparison of AXL expression in YFP^-^ and YFP^+^ (B-pDCs). The results are representative of two independent experiments each containing at least 3 mice per group. Mann-Whitney tests were used as statistical method.

**Supplemental Figure 5.**
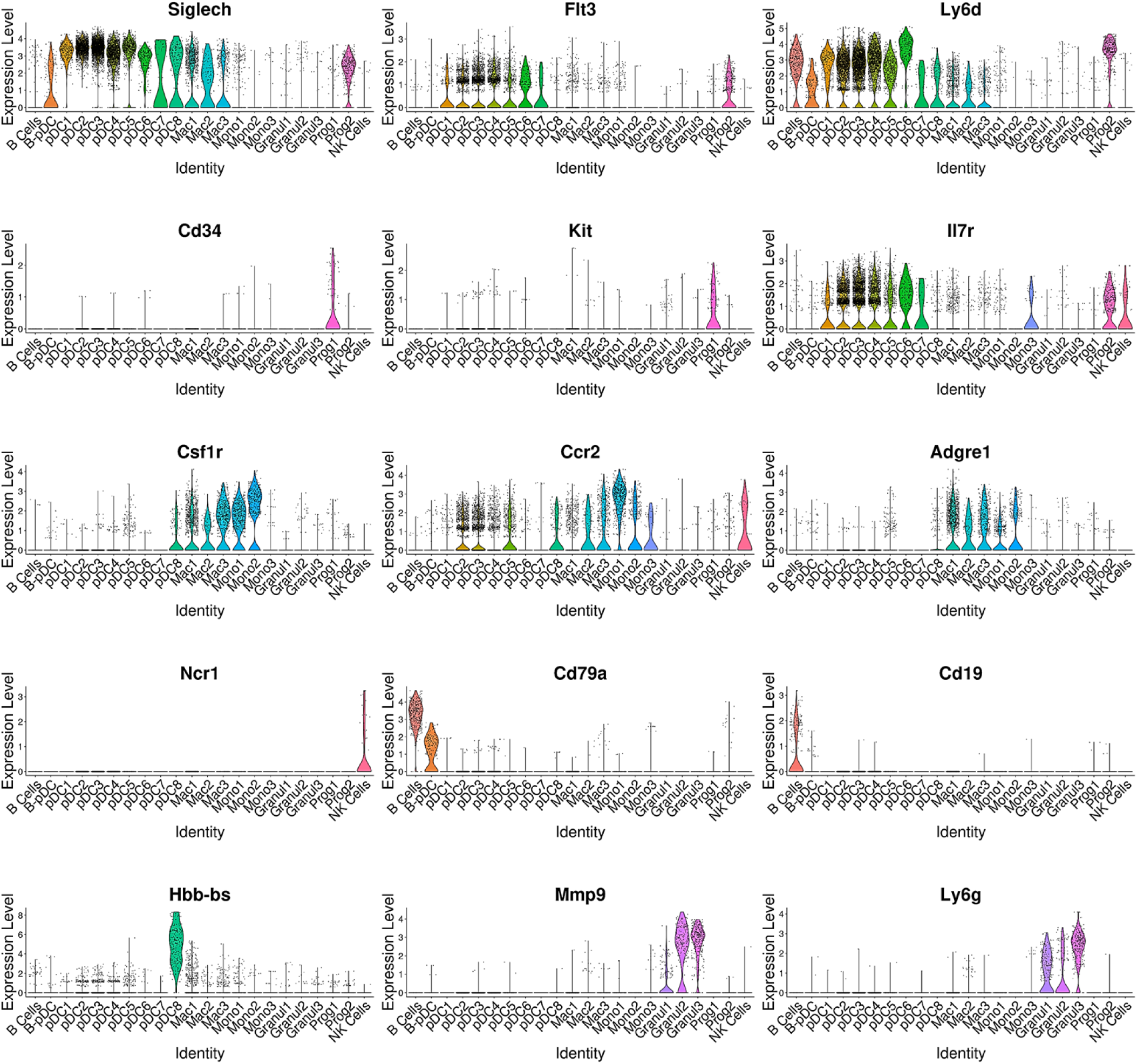
Confirmation of unbiasedly identified cell cluster identities. Seurat generated violin plots of genes associated with distinct cell populations including mature B cells, pDCs, granulocytes, pre-pDCs, NK cells, monocytes, macrophages, and myeloid progenitor cells. Plot identities were initially determined by running the package CIPR (Cluster Identity Predictor).

**Supplemental Figure 6.**
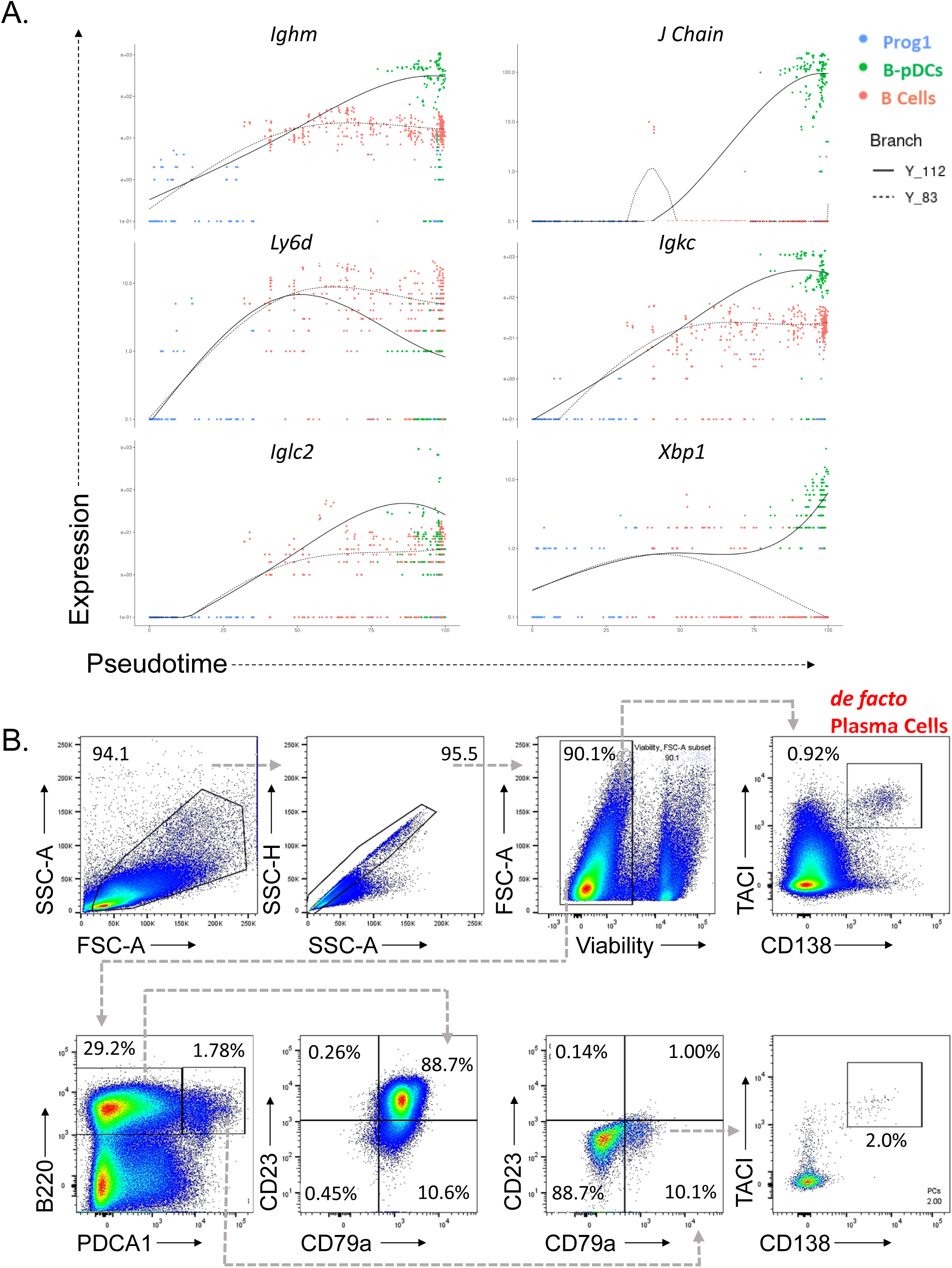
Expression of early B cell receptor genes in B-pDCs. (A) Monocle was used to plot select early B cell genes as function of branched pseudotime for Prog1, B Cells, and B-pDC populations. (B) Plasma cell contamination in *Mb1*^+^ pDCs (B220^+^ PDCA1^+^) was ruled out by flow cytometric analysis of TACI and CD138 in BM of 6 C57BL/6J mice.

**Supplemental Figure 7:**
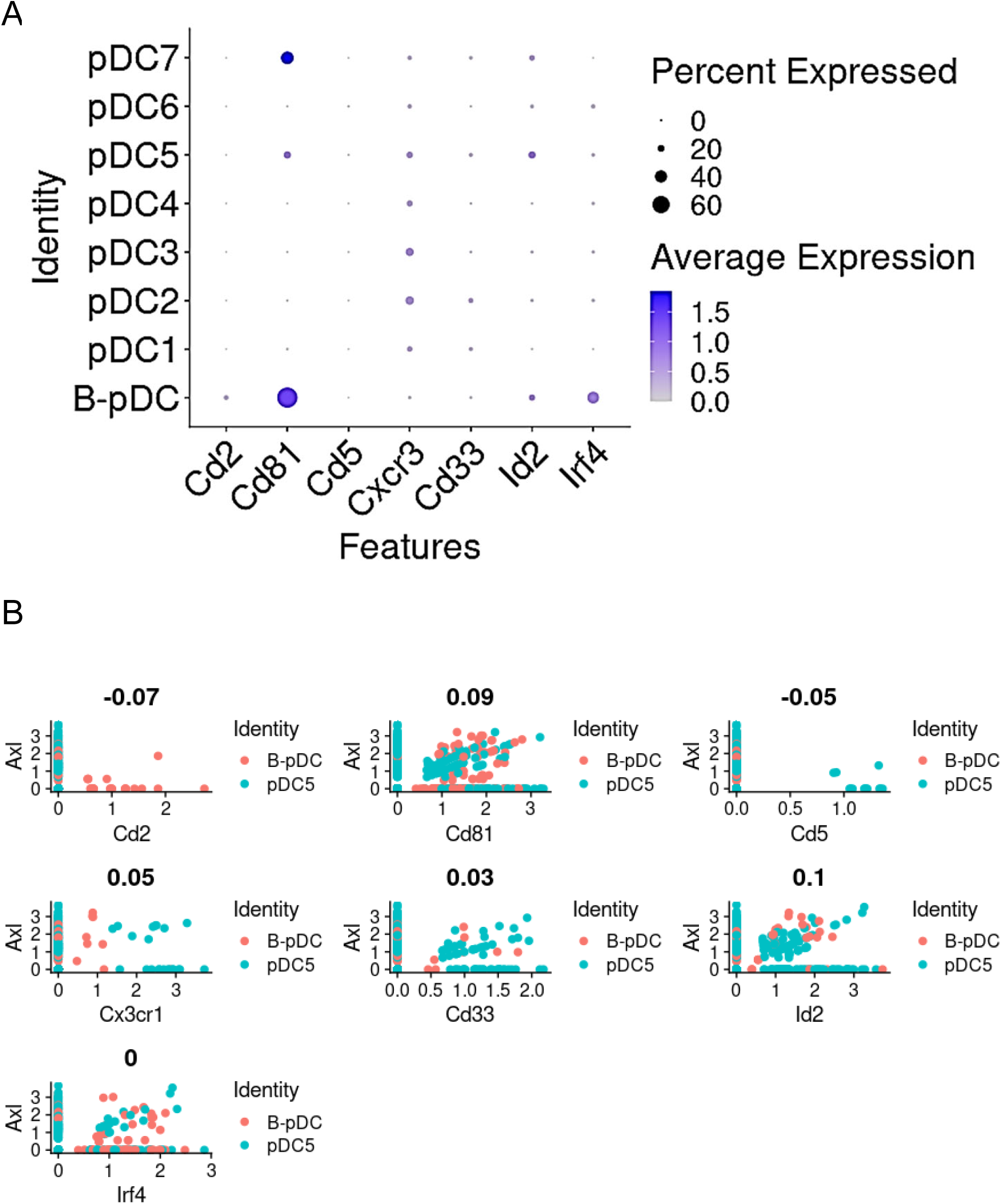
Transcriptional heterogeneity of *Axl^+^* murine pDC populations. (A) Dot plot depicting expression prevalence of select “*AXL^+^* DC” associated markers from literature select and pDC DEGs for pDC subsets. (B) Correlation between expression of AXL+ DC associated markers and *Axl* in non-canonical mouse pDC subsets.

**Supplemental Table 1.**
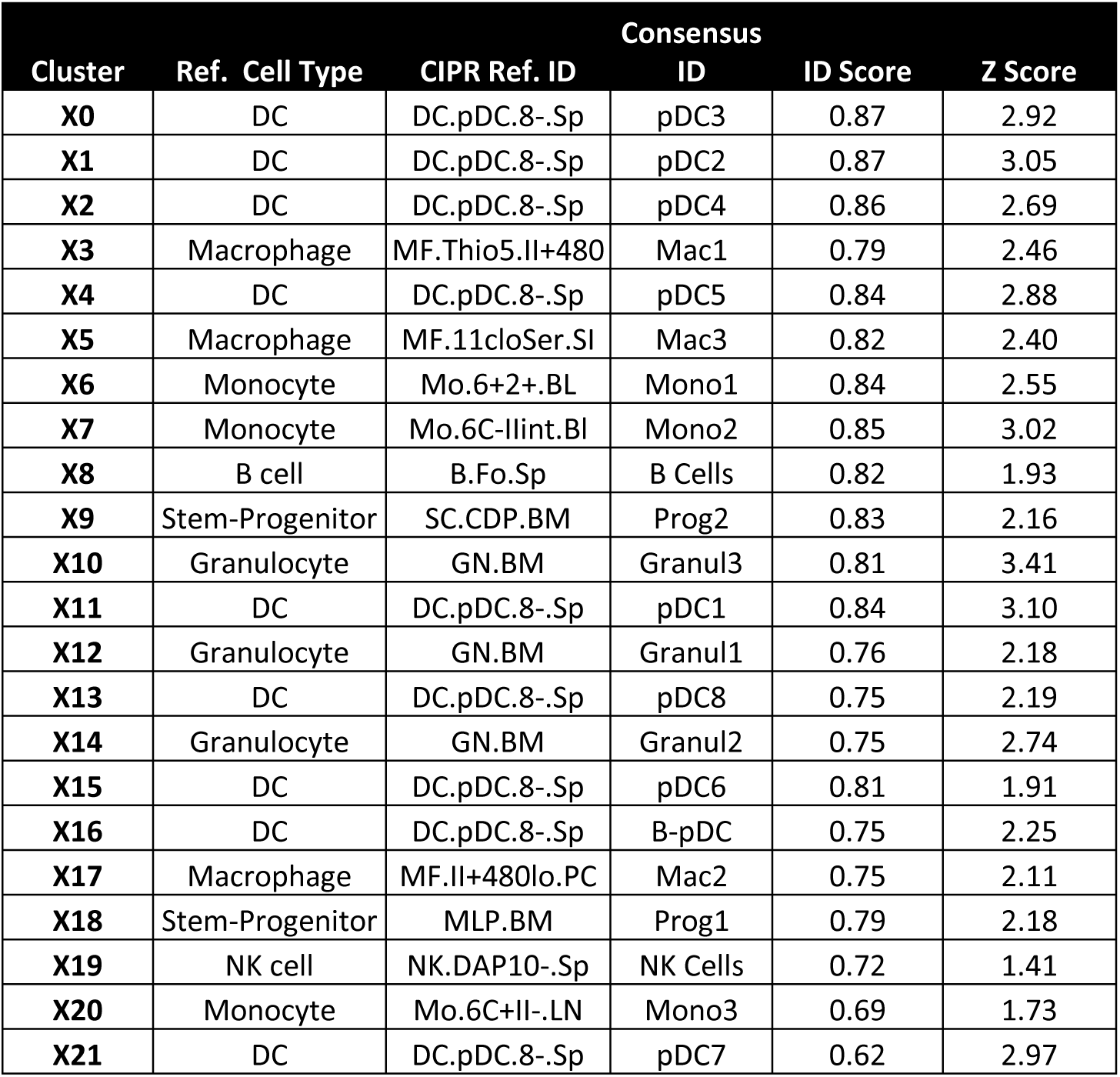
Top CIPR Assigned Identity Scores for Seurat Cluster DEGs. CIPR IDs were generated using the top 1000 DEGS from each cluster. The top 5 consensus CIPR ID generated was used to rename Seurat clusters.

**Supplemental Table 2.**
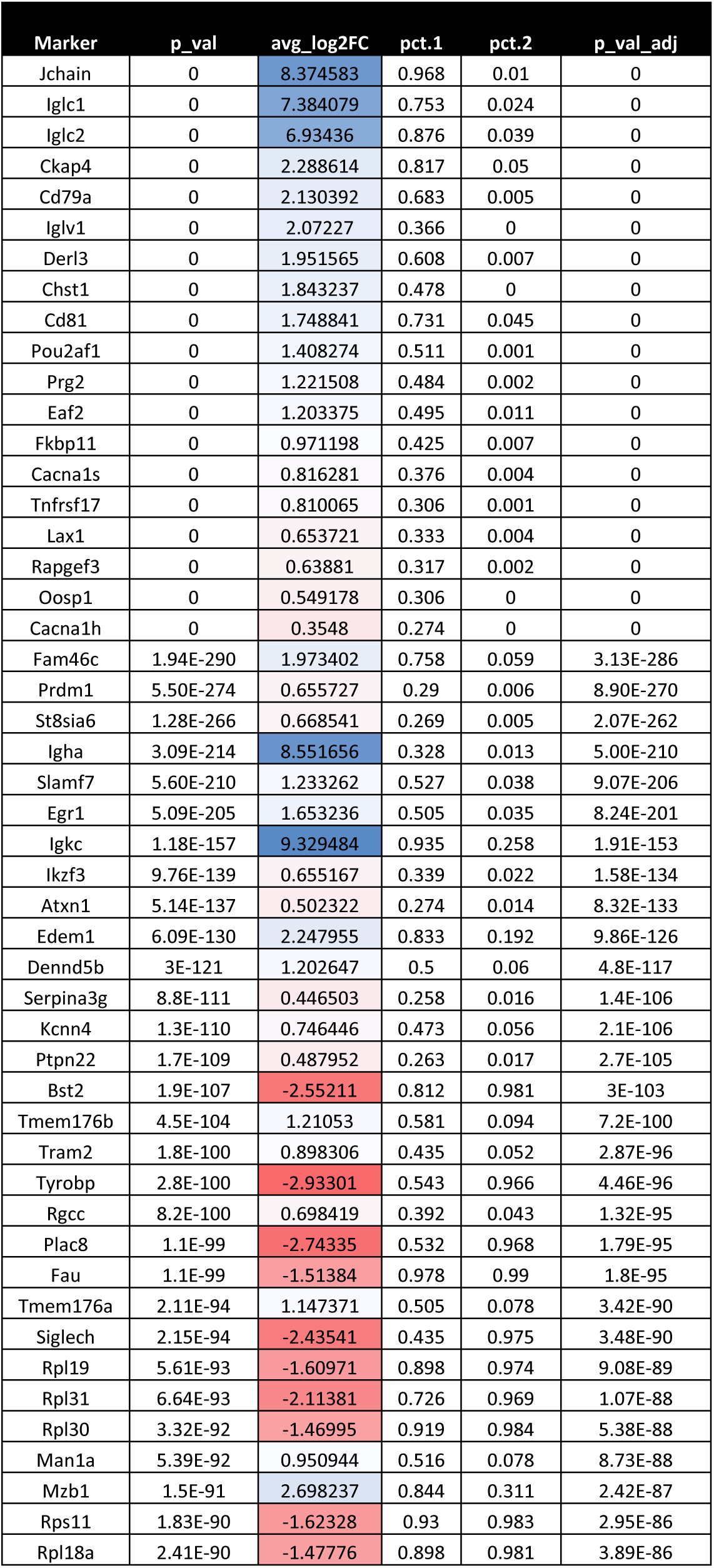
DEG comparison between progenitor clusters. The top 50 differentially expressed features of Prog1 in relation to Prog2 were calculated using Seurat using non-parametric Wilcoxon rank sum test. In blue are upregulated genes and in red are downregulated genes.

## Methods

### Mice

Generation of *Bcl11a* cKO Mice (C57BL/6J background) was performed as described^14^. To generate conditional knockouts for the *Mb1* gene, specifically, *Bcl11a* ^F/F^ mice were crossed to *Mb1*-Cre^+^ Rosa26-YFP^+^ reporter mice. *Mb1*-Cre deleter and Rosa26-YFP reporter strains (C57BL/6J genetic background) were obtained from Jackson labs (catalog #’s 020505 and 006148, respectively), bred, and genotyped according to the vendor protocol and their validated primers). All housing, husbandry, and experimental procedures with mice were approved by the Institutional Animal Care and Use Committees at the University of Texas at Austin. 6-week-old gender matched male or female mice were used for all experiments described in this manuscript. For each experiment, at least 4 samples per experimental group were used, with each experiment repeated at least 3 times (except for bone marrow reconstitutions and imaging flow cytometry experiments, which were repeated twice).

### Tissue processing

Mouse femurs were cut at the extremities and a 25G needle was inserted in the bones to flush out bone marrow cells with 10 ml of cold PBS onto a 70 uM strainer in a 50 mL tube. Cells were washed from the strainers into the tubes with an additional 10 mL of PBS. Tubes were spun at 300G for 5 minutes and supernatant was decanted. 2 mL of ACK red blood cell lysis buffer was used to resuspend cells, which were incubated for 2 minutes and 20 seconds. Samples were diluted with 20 mL of cold PBS, washed once, re-filtered and resuspended in 200 uL of FACS buffer for subsequential flow cytometry staining. Spleens were mashed onto pre-wet 70 uM filters in a 50 mL tube with syringe plungers and 20 mL of cold PBS was added through the strainers into the tube. Cells were spun at 300G for 5 minutes and supernatant was decanted. 2 mL of ACK red blood cell lysis buffer was used to resuspend cells, which were incubated for 2 minutes and 20 seconds. Samples were diluted with 20 mL of cold PBS, washed once, re-filtered and resuspended in 200 uL of FACS buffer for subsequential flow cytometry staining.

### Flow cytometry

Single cell suspensions (1x10^6^ cells) were resuspended in 200 uL of FACS buffer (5mM EDTA and 1% Heat Inactivated Fetal Calf Serum in D-PBS) and incubated for 15 minutes with surface antibody cocktail (each antibody at a concentration of 1.25 micrograms/ml in FACS buffer) in the dark at 4C. Cells were then washed 3 times with FACS buffer and ran live on a BD LSRII Fortessa. Single color controls for panel compensation were made using AbC Total Antibody Compensation Bead kit (ThermoFisher Scientific), by staining each control with one of the panel antibodies and using these controls to compensate the flow cytometer experiment prior to running the samples. A non-stained sample of cells was included in experimental runs and used to adjust for autofluorescence and any nonspecific staining. FMO single cell controls were also included in some of the experimental runs (cells stained with all antibodies in the panel minus one), for compensation quality control and to detect positive MFI thresholds for each fluorophore using cells. Sorting was performed on a FACS Aria (BD Biosciences). All analysis was done using FlowJo (Tree Star) software. Single-cell suspensions were stained with the following antigen-specific monoclonal antibodies (clones in parenthesis): anti CD45R/B220-BV605 (RA3-6B2), CD19-Alexa Fluor 700 (6D5), CD86-APC-Cy7 (GL-1), I-A/I-E-Pacific Blue (M5/114.15.2), IL7R-BV421 (A7R34), cKit-PE-Cy7 (ACK2), CD11b-PerCP-Cy5.5 (M1/70), CD3e-PerCP-Cy5.5 (500A2), Gr1-PerCP-Cy5.5 (RB6-8C5), CD4-PerCP-Cy5.5 (RM4-5), CD8-PerCpCy5.5(53-6.7), NK1.1-PerCp-Cy5.5 (PK136), Sca1-BV711 (D7), CD135-APC (A2F10), CD150-BV605 (TC15-12F12.2), CD79a-FITC (24C2.5) (Invitrogen), Anti-CD11c-PerCP-Cy5.5 (N418) (eBioscience), Anti-AXL-APC (175128)(R&D Systems), Anti-CD23 BV510 (B3B4) (Biolegend), Anti-CD267-BV421 (8F10) (BD Biosciences), Anti-CD138-APC (281-2) (Biolegend), Anti-CD83-PeCy7 (Michel-19), and Anti-PDCA1-PE or Biotin (JF05-1C2.4.1) (Miltenyi Biotech), in D-PBS/2% (vol/vol) FBS FACS buffer. In some experiments, viability stain Ghost Dye Red 780 (Fisher Scientific) was used to stain cells in PBS prior to antibody staining according to the manufacturer ‘s protocol. Flow cytometry data was analyzed with FlowJo, while Imaging flow cytometry was performed on ImageStream X and analyzed on IDEAS® 6.0 (Amnis), by pre-filtering out of focus cells and performing automatic software guided compensation with single color controls.

### Bone Marrow Transplantation

Bone marrow from 6-week-old *Bcl11a*^F/F^/mb1-Cre ^+^ cKO or Mb-1-Cre-YFP reporter mice (n=3 per group) were collected from femurs and 5x10^6 bone marrow cells were transferred via retro-orbital injection into recipient immunocompetent C57BL/6J mice lethally irradiated with two doses of 450 rad 1 hour apart. Mice were kept on an antibiotic diet for 3 weeks to allow for immune reconstitution. Eight weeks post transfer, mice were sacrificed and cells collected from BM and spleen to investigate reconstitution of cellular subsets.

### RNA isolation and RNA-seq

For sorting prior to RNA collection, BM from 12 mice was prepared from femurs at 6 weeks of age, combined into three groups (4 mice/group). Total RNA was extracted from FACS sorted *Bcl11a* cKO BM pDCs (approximately 100K YFP^+^ pDCs and 400K YFP^-^ pDCs per pooled sample) using TRIzol reagent (Invitrogen). Taq polymerase (New England Biolabs) and a Perkin-Elmer 2700 thermocycler were used to amplify transcripts for the following mouse genes: *Bcl11a* (F: 5’-GTGGATAAGCCGCCTTCCCCTT-3’, R: 5’- GGGGACTTCCGTGTTCACTTTC-3’), *Bst2* (F:5’- AGGCAAACTCCTGCAACCTG-3’, R: 5’- ACCTGCACTGTGCTAGAAGTC-3’), *SpiC* (F: 5’- ATCCTCACGTCAGAGGCAAC-3’, R: 5’-TGTACGGATTGGTGGAAGCC-3’, *Id2* (F: 5’- GGACATCAGCATCCTGTCCTTGC- 3’ R: 5’- GTGTTCTCCTGGTGAAATGGCTGA- 3’,and *β-actin* (F:5’AGGTGTGATGGTGGGAAT-3’, R: 5’- GGTGTAAAACGCAGCTCAGT-3’) and probe RNA sequencing potential. For bulk RNA-seq, twelve mice were pooled into three YFP^+^ groups, and total RNA was extracted as above. Oligo-dT-primed cDNA was prepared using SuperScript III First-Strand Synthesis System for RT-PCR (Invitrogen). Poly(A) mRNA was enriched using the NEBNext magnetic isolation module (E7490) and samples underwent DNAse treatment. cDNA was prepared using the Ultra Low kit from Clontech (Mountain View, CA). Libraries were prepared at the according to manufacturer’s instructions for the NEBNext Ultra II Direction RNA kit (NEB, product number E7760). The resulting libraries tagged with unique dual indices were checked for size and quality using the Agilent High Sensitivity DNA Kit (Agilent). Library concentrations were measured using the KAPA SYBR Fast qPCR kit and loaded for sequencing on the Illumina NovaSeq 6000 instrument (paired-end 2X150, or single-end, 100 cycles; 30x10^6 reads/sample). Data were analyzed using a high-throughput next-generation sequencing analysis pipeline: FASTQ files were aligned to the mouse genome (mm9, NCBI Build 37) using TopHat2^53^. Gene expression profiles for the individual samples were calculated with Cufflinks54 as RPKM values. YFP^+^ pDC samples were normalized to each control YFP-pDC sample. GO terms identified as significantly different: GO:0006955, GO:0009611, GO:0048584, GO:0006954, GO:0002684, GO:0001775, GO:0009986. Panther terms: BP00148, BP00155, BP00102, BP00120. Gene Set Enrichment Analysis (GSEA) using ordered gene expression levels of Common Lymphoid Progenitor-derived B-pDC and Common Myeloid Progenitor-derived pDC were significantly enriched in both GO terms and Panther-derived gene sets. A randomly selected control GSEA curated set, GSE7831: UNSTIM_VS_ INFLUENZA_STIM_ PDC_4H_DN (defined as genes down-regulated in untreated pDC versus influenza virus infected pDC 55 showed insignificant enrichment. Normalized enrichment score (NES) and false discovery rate q-values (FDR); FDR ≤ 0.25 is considered significant^56^.

### TLR9 engagement

Mb1-cre-YFP reporter mice were injected via tail veins with 100ul of PBS containing 50ug of CpG:ODN (n=4) or GpC:ODN control (n=4) (CpG-B no.1826, TCCAT GACG TTCCT GACGTT; control non-CpG-B no.2138, TCCATGAGCTTCCTGAGCTT, Invivogen, USA). At 24 hours, mice were sacrificed and bone marrow and spleens were collected for pDC phenotyping.

### Cell co-cultures

Mb1-YFP^+^ B-pDC (B220^+^ PDCA1^+^YFP^+^) and Mb1-YFP^-^ pDCs (B220^+^PDCA1^+^YFP^-^) subsets were sorted on a FACS Aria (BD Biosciences) by and cultured in 96-well round bottom plates (n=3 wells in triplicates, repeated twice) with RPMI medium 1640 containing 10% (vol/vol) FBS, 2 mM L-glutamine, 100 units/mL penicillin and streptomycin, 1 mM sodium pyruvate, and 10 mM Hepes, and with or without 5 μg/ml class A CpGs (ODN 1585, InvivoGen) for 24 hours. pDCs were then washed three times to remove residual CpG. Preactivated pDCs (5 × 10^3^) were cultured with lymphocytes (2.5 × 10^4^) magnetically sorted frommouse spleen single cell preparations using MACs columns and the EasySep™ Mouse T Cell Isolation Kit (StemCell Technologies Cat# 19851) for a total of 6d with 20 units/mL of IL-2. CSFE labelling of lymphocytes (for which sorting purity was determined to be 97% via flow cytometry) prior to coculturing experiment was done using the CellTrace™ CFSE Cell Proliferation Kit (ThermoFisher Scientific; Cat# C34554) accordingly the manufacturer’s protocol.

### Enzyme-linked immunosorbent assays (ELISA)

Sorted B-pDC and pDCs (5 × 10^3^ sorted cells per well, n= 3 per group, done in duplicate wells for each condition) were stimulated with 5 uM CpG-A (ODN 1585, Invivogen) or CpG-C (ODN 2395, Invivogen); or left untreated for 24 hs. ELISAs for IFN-α (Invitrogen) or IL-12p40 (BioLegend), respectively, was performed in 100 uL of cell free supernatants according to manufacturer’s instructions. Data was acquired using a Bio-Tek model EL311 automated microplate reader measuring absorbance at 450 nm.

### Statistical analysis

All flow cytometry or ELISA data were analyzed with Prism (Version 8; GraphPad, La Jolla, CA) using Mann-Whitney non-parametric t-tests. All bar graphs show means and standard deviation (SD) and are representative of repeated experiments. These data were considered statistically significant when p-values were <0.05.

### 10x Single cell RNA-seq analysis

PDCA1^+^ C57BL/6J bone marrow cells from 4 pooled 6 week old wildtype C57BL/6J mice were magnetically sorted using MACS columns into culture medium, washed once with PBS^+^ 0.04% BSA, and re-suspended in 32 μl PBS ^+^ 0.04% BSA. Single cell suspensions (50,000) were processed in the University of Texas Genomic Sequencing and Analysis Facility (GSAF). Cell suspensions were loaded on the Chromium Controller (10X Genomics) and processed for cDNA library generation following the manufacturer’s instructions for the Chromium NextGEM Single Cell 3’ Reagent Kit v3.1 (10X Genomics). The resulting libraries were examined for size and quality using the Bioanalyzer High Sensitivity DNA Kit (Agilent) and their concentrations were measured using the KAPA SYBR Fast qPCR kit (Roche). Samples were sequenced on the NovaSeq 6000 instrument (paired end, read 1: 28 cycles, read 2: 90 cycles) with a targeted depth of 71,130 reads/cell. Cellranger (v3.0.2) was used to demultiplex samples, to map data to the mouse transcriptome (mm10) and quantify genes. The gene counts matrix was read into Seurat (v3.2.1). Cells with unique feature counts over 3,500 or less than 200, and >3% mitochondrial read counts were removed from the analysis. The data was normalized and transformed using scTransform. Cells were clustered based on the top 1000 variable features, 20 PCs, and a resolution of 0.6 (Seurat’s graph-based clustering approach), and then visualized using UMAP. Analysis of the normalized and filtered single cell gene expression data (median of 16,181 genes across 13,838 single-cell transcriptomes) was used for the final expression table and downstream analysis. Wilcoxon rank sum test was used to test for differential expression among clusters and identify gene markers (logfc.threshold = - 1, min.pct= 0.25). Gene markers and other genes of interest were visualized using violin plots, dot plots, and heatmap plots. Unsupervised ordering of PDCA1^+^ cells was done with the Seurat integrated results as input to build a tree-like differentiation trajectory using the DDRTree algorithm of the R package Monocle v2 ^59^. Select DEGs of B-pDCs, B Cells, and Prog1 populations were set as ordering genes for each trajectory, with the root state set as Prog1 classified cells. Expression of genes as a function of pseudotime was plotted with the plot_genes_branched_pseudotime Monocle function. We also used Monocle3 to generate partition-based learning of single cell developmental trajectories with SimplePPT and plot UMAP based pseudotime figures with initial reclustering parameters num_dim=60, resolution=1e-5.

### Data Deposition

RNA-seq: GSE105827; ChIP-seq: GSE99019; 10x ScRNA-seq: GSE225768. Accession numbers to previously published data sets: GSE52868 (pre-B RNA-seq); GSE55043 (CAL1 ChIP-seq); ENCODE BCL11A ChIP-seq in GM1287838.

## Author contributions

J.D.D. and G.C.I. designed research; A.A., J.D.D., K.G., C.R., Z.H., B-K.L., and G.C.I. performed research; J.D.D., J.L. V.R.I, L.I.E., S.Y., G.G., A.A., and G.C.I., contributed new reagents/analytic tools; A.A., J.D.D., C.R., B.-K.L., S.Z., D.O.H., and G.C.I., analyzed data and A.A. and J.D.D. wrote the paper. J.D.D is solely responsible for all flow cytometric experiments/analysis and A.A. is responsible for the bioinformatics portion. The authors declare no conflict of interest.

## Acknowledgements

We thank June V. Harriss for expert assistance in the generation of *Bcl11a* conditional knockout mice, and Chhaya Das and Maya Ghosh for help in ChIP experiments and cell culture. The CAL1 cell line was kindly provided by Drs. Takahiro Maeda and Boris Reizis. Library preparation and Illumina ChIP- and RNA-seq were performed at the NGS core of the MD Anderson Cancer Center. Single Cell RNA-seq was performed at the University of Texas Genomic Sequencing and Analysis Facility with the help of Holly Stevenson and Dhivya Arasappan. The Lymphoma Research Foundation Fellowship 300463 (to J.D.D.); NIH grant F32CA110624 and Owens Medical Research Foundation grant (to G.C.I.); NIH grant R35GM133658 (to S.Y.); NIH grant R01AI104870 (to L.I.R.E.); NIH grant R01CA130075 (to V.R.I.); NIH Grant R01CA31534; Cancer Prevention Research Institute of Texas (CPRIT) Grant RP120348 to the MD Anderson NGS core and CPRIT RP120459 and the Marie Betzner Morrow Centennial Endowment (to H.O.T.) provided support for this work.

